# Sensitivity to Error During Visuomotor Adaptation is Similarly Modulated by Abrupt, Gradual and Random Perturbation Schedules

**DOI:** 10.1101/2021.06.14.448375

**Authors:** Susan K. Coltman, Robert J. van Beers, W. Pieter Medendorp, Paul L. Gribble

## Abstract

It has been suggested that sensorimotor adaptation involves at least two processes (i.e., fast and slow) that differ in retention and error sensitivity. Previous work has shown that repeated exposure to an abrupt force field perturbation results in greater error sensitivity for both the fast and slow processes. While this implies that the faster relearning is associated with increased error sensitivity, it remains unclear what aspects of prior experience modulate error sensitivity. In the present study, we manipulated the initial training using different perturbation schedules, thought to differentially affect fast and slow learning processes based on error magnitude, and then observed what effect prior learning had on subsequent adaptation. During initial training of a visuomotor rotation task, we exposed three groups of participants to either an abrupt, a gradual, or a random perturbation schedule. During a testing session, all three groups were subsequently exposed to an abrupt perturbation schedule. Comparing the two sessions of the control group who experienced repetition of the same perturbation, we found an increased error sensitivity for both processes. We found that the error sensitivity was increased for both the fast and slow processes, with no reliable changes in the retention, for both the gradual and structural learning groups when compared to the first session of the control group. We discuss the findings in the context of how fast and slow learning processes respond to a history of errors.

**New & Noteworthy:** We investigated what aspects of prior experience modulate error sensitivity, within the framework of a two-state model of short-term sensorimotor adaptation. We manipulated initial training on a visuomotor adaptation reaching task using specific perturbation schedules that are thought to differentially affect fast and slow learning processes, and we tested what effect these had on subsequent adaptation. We found that sensitivity to adaptation error was similarly modulated by abrupt, gradual, and random perturbation schedules.

## Introduction

Adaptation is often defined as an error-driven process, in which the error experienced during a movement leads to a corrective adjustment in the motor output on the following movement (1-5). Behavioural measures of adaptation are well characterized by state-space models (1, 4), which represent trial-to-trial changes in movement as a function of how an error on a given trial affects motor output on the subsequent trial. The update from one trial to the next, or the change in motor output, is based on two parameters: a retention parameter which determines what proportion of motor output is retained from trial to trial, and an error sensitivity parameter which governs the proportion of error experienced on the current trial that is corrected for on the subsequent trial.

Variations of the state-space model are built on the assumption that adaptation is the product of multiple underlying processes with distinct timescales (3, 6-8). Researchers have begun to provide neural evidence to strengthen the theory that sensorimotor learning is supported by multiple processes (9, 10). An influential two-state model of short-term motor adaptation was proposed by Smith et al. (3) that proposed a fast process that learns quickly but has poor retention and a slow process that learns more slowly, but has strong retention.

The prevailing success of the two-state model continues to be that it accounts for the learning phenomenon known as savings, characterized as prior learning speeding up subsequent relearning (3, 11). While Smith et al. (3) initially argued that the reason for the fast relearning during a second introduction of the same perturbation was due to the resistance of the slow process to change, recent studies suggest that learning rate can be modified depending on factors such as the uncertainty of movement error (12, 13), size of movement error (14), and a history of movement errors (15-17).

The behavioural changes associated with savings suggest that some component of memory from the initial training must lead to the faster relearning, but what is remembered and recalled remains unclear (15-22). One perspective argues for the enhancement of an explicit strategy (18, 20, 23), while the other side suggests that faster relearning is driven by the experience of the motor errors (15-17, 24).

In support of the latter possibility, Herzfeld and colleagues (16) proposed that a history of errors modulates the error sensitivity on each trial, systematically controlling how much the motor system learns from the current motor error. They suggested that an error-based adaptation model that provides for experience-dependent error sensitivity modification could account for savings. Furthermore, Leow et al. (17) demonstrated that it is a memory of errors, not previous actions, that is necessary for savings.

Recent work has shown that repeated exposure to the same force field perturbation results in greater error sensitivity of both the fast and slow processes (15). While in Coltman et al. (15), the error sensitivity terms for the fast and slow processes were held constant within a session, we evaluated the theory of experience-dependent error sensitivity modulation in the context of changes in error sensitivity from one learning session to the next. Although these results clearly indicate that the motor system stored some component (i.e., memory) of prior training to speed up subsequent learning, it remains unclear how the fast and slow learning processes contribute to savings and what aspects of prior experience modulate error sensitivity. In other words, do both fast and slow processes access a single stored component of prior training, or do they store independent components? If the fast and slow processes depend on separate stored components of prior training, then it may be possible to independently modulate characteristics of the fast and slow processes (e.g., error sensitivity) by experimentally manipulating aspects of prior learning.

Recent findings demonstrate that participants do not adapt linearly in response to different magnitudes of error (13, 14, 25). Additionally, Orban de Xivry and Lefevre (26) propose that different perturbation schedules lead to distinct motor memories with different attributes and neural representations (i.e., the amount of reorganization of the motor cortex). We propose that perturbation schedules that are designed to produce learning using errors of different magnitudes may have a differential effect on session-to-session changes to the fast versus slow processes. In the present study, we manipulated initial training in a visuomotor adaptation task using perturbation schedules which involved errors of different magnitudes, and we tested what effect these different initial learning experiences had on subsequent adaptation, and specifically on characteristics of the fast versus slow adaptation processes.

We asked one group of participants to counter a gradual perturbation schedule during initial training. When a perturbation is gradually introduced, such that participants never experience large errors, learning is believed to be more implicit in nature (26). We predicted that when participants in this group were later tested on an abrupt perturbation, only the slow process would be affected by the initial training, compared to a control group who were initially trained using an abrupt perturbation. For a second group of participants, initial training was based on a structural learning paradigm, involving a series of brief exposures to large, random perturbations (27, 28). This perturbation schedule is thought to be based on explicit learning mechanisms (20, 29). During the original conception of the two-state model, Smith et al. (3) designed a rebound paradigm to test the theorized characteristics of each process. Much like the brief reversal used in the rebound paradigm, which includes large errors and limited exposure, the motor output during this phase of the paradigm is believed to be dependent on the fast process (3). For this group we predicted that when later tested on an abrupt perturbation, only the fast process would be affected by the initial training, as compared to the control group.

We modelled perturbation-driven changes in movement with the state-space equations proposed by Smith et al. (3), and focused on changes in the retention and error sensitivity parameters. The model estimates function as a tool for understanding how the underlying processes of adaptation were affected by the prior training. Substantiating the finding of Coltman et al. (15), we confirm that repetition of the same visuomotor perturbation results in an increase in error sensitivity for both processes, when comparing the two sessions of the control group. By comparing the model estimates of participants in the gradual and structural learning groups to the first session of the control, we expected to see changes in error sensitivity that depended on the type of prior training participants experienced. Interestingly, however, we found that error sensitivity of both the fast and slow processes was increased for both groups. The findings are discussed in the context of storing and accessing a history of errors.

## Methods

### Participants

A total of 60 healthy young adults (age range 21-35; mean age ± sd 27.9 ± 4.2 years) participated in a visuomotor rotation experiment. Participants were recruited from the online platform maintained by Prolific.co and received £11.25 for their participation. As part of the Prolific platform, participants respond to a series of questions related to age, gender, health and economic status. Based on this prescreen information, 24 participants identified as female and 36 as male. Participants were recruited globally and reported being located in 17 different countries (Estonia, Finland, France, Greece, Hungary, Israel, Italy, Mexico, Netherlands, Poland, Portugal, Slovenia, South Africa, Spain, Sweden, United Kingdom and the United States). All participants self-reported being right-handed and had normal or corrected-to-normal vision. The protocol was approved by Western University’s Research Ethics Board and all participants indicated electronic consent.

### Apparatus

Participants used a standard computer mouse and their own computer to access a webpage hosted on a network computer located at Western Interdisciplinary Research Building. The task was written in and controlled by JavaScript, running locally within the participants’ web browser.

Participants were asked to use a standard computer mouse and a standard credit or debit card to complete a spatial calibration procedure. Participants were initially instructed how to turn off the acceleration for the mouse, based on their operating system. Then, following an instruction video, participants were asked to align the top of their mouse with the top of the credit card. After a tone, they were instructed to move the mouse in a smooth and straight path, aligning the top of their mouse with the bottom of the card. Participants were asked to hold still while waiting for a second tone, indicating that they needed to realign the mouse with the top of the card. This was repeated at two different speeds indicated in the video. When the calibration procedure was successfully completed, participants watched an instructional video about the experimental task.

The size and position of the stimuli were scaled based on a mouse calibration procedure. Real-time position of the mouse was used to control the visual display and to provide on-line visual feedback. The mouse speed was adjusted such that the distance from start position to target was exactly 6 cm based on the calibration. While the physical target distance was always 6 cm, this translated to 300 pixels on screen. Therefore, the straight reach trajectory was 300 pixels, however a participant’s view of this was potentially compressed or expanded relative to the target value of 6 cm, depending on their monitor as well their viewing distance from the monitor.

### Paradigm

At the start of each trial, participants were instructed to click their mouse to begin. A circular cursor (10 pixels radius) was virtually displayed on the participant’s computer monitor and was used to represent the position of the mouse on screen. The position of the mouse at the start of the trial, represented the start position on screen. A small square (20 pixels by 20 pixels) represented the target. The radial distance of the target from the start position was 300 pixels. The target appeared at either 45°, 90°, or 135°, relative to the start position (where 6 cm directly to the right of the start position represented 0°). The location of the target was randomized per trial, per participant, such that each participant saw a different order of targets with an equal number of presentations of each target over the course of a session.

Participants were instructed to make a straight movement from the start position to the target, within a narrow temporal window. At the beginning of each trial the target appeared in white. Participants were required to hold still at the start position for 500 ms, at which time the target changed color to green, representing a “go” signal for participants to initiate a movement to the target. In addition to the colour change of the target, a tone was used as a secondary “go” signal. Participants needed to reach for the target and bring the centre of a red cursor representing the position of their computer mouse within 10 pixels of the centre of the target within 600–900 ms. If a participant’s movement time was less than 600 ms, the target turned red to indicate that the movement was “too fast”. If the participant’s movement time was within 600–900 ms, the target remained green to indicate that the movement was “good”. If the participant’s movement time was greater than 900 ms, the target would turn blue to indicate “too slow”. Feedback related to movement time was displayed on the screen for 1000 ms before the screen went blank and written instructions on screen indicated that the participant should return the mouse to a comfortable starting position within their workspace. Participants were instructed to try to obtain the “good” feedback as often as possible throughout the experiment.

To assist with making straight movements between the start position and the target using a computer mouse, the first 20 trials of the first session represented a practice session for participants. In these trials, a purple rectangle (50 pixels by 300 pixels), with two white lines on either side was shown on screen, highlighting a straight path to the target. Participants were instructed to keep the red cursor on the path, between the lines, toward the target. If the cursor moved outside the path, the background colour changed from black to pink.

Participants were randomly assigned to one of three groups. Each group completed two sessions (initial training and testing), separated by a 5-minute break (**Fig. 1**). Each session included a total of 450 reaching movements, with a 1-minute mid-session break halfway. The experimental paradigm for each session consisted of 4 epochs. The first epoch (baseline) consisted of 70 trials in which participants were provided with veridical feedback of the cursor position. The second epoch (adaptation) consisted of 300 trials in which a visuomotor rotation was applied to the cursor feedback: an angular rotation was imposed on the cursor, such that a hand movement aimed directly at a target produced a cursor movement that was rotated radially about the start position and participants saw that their movement had generated an error. Participants had to learn to counter the rotation by moving their hand in an equal and opposite direction. With practice, participants adjusted their movements in such a manner that the visual feedback produced straight trajectories from start position to the target. In the third epoch (error-clamp; consisting of 30 trials) the task error was clamped to zero. During the clamp trials, the angular position of the cursor relative to the start position was clamped to a straight line connecting the start position to the target, while participants maintained control of the radial distance of the cursor from the start position. Finally, in the fourth epoch (washout; consisting of 50 trials), participants were provided again with veridical feedback to bring performance back to baseline.

**Figure 1.**
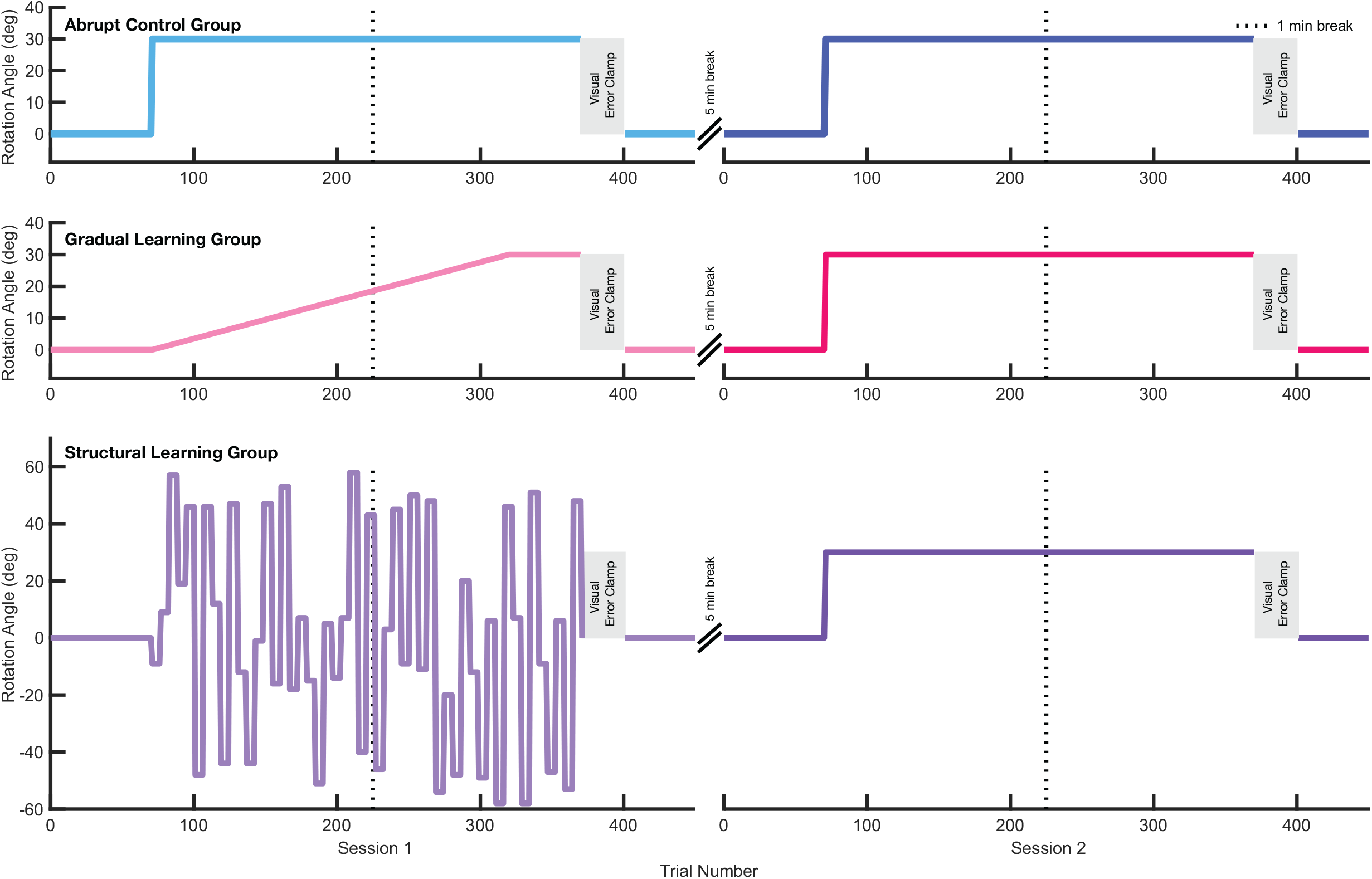
Experimental design and perturbation schedule. The experiment was divided into two sessions, separated by a 5-min session break. Each session consisted of four blocks: 1) a baseline period of no rotation trials, 2) an adaptation period, 3) an error clamp period, and 4) a washout period. Participants were randomly assigned to one of three groups which differed in session one during the adaptation period: abrupt control group, gradual learning group, or structural learning group.

During the adaptation epoch of the first session, participants experienced one of three conditions: (1) a control learning group (n=20) experienced an abrupt 30° clockwise (CW) rotation for all 300 trials during this phase (**Fig. 1; top**), (2) a gradual learning group, (n=20) in which a rotation was increased linearly from 0° to 30° CW over 250 trials and then held at a fixed 30° CW for another 50 trials (**Fig. 1; middle**), or (3) a structural learning group (n=20) in which participants encountered random rotations, ranging from 60° counter-clockwise (CCW) to 60° CW in blocks of 6 trials with the same rotation (27, 28, 29; Fig. 1; bottom). In this group, we deliberately set the average over all angles to zero, to prevent any accumulative learning. We also excluded rotation sizes within 10° of the test rotation (30° CW) and its inverse (30° CCW). We furthermore set the change in rotation angle to be equal to or greater than 15° to ensure the errors were always large, which characteristically has the greatest influence on the fast process (3, 29). During the second session, all three groups experienced an abrupt 30° CW rotation during the adaptation epoch.

### Data Analysis

The position of the cursor in both x (lateral) and y (sagittal), were sampled in pixels at the refresh rate of their computer monitor (typically 60 Hz). Missed samples were interpolated during analysis (less than 1 % of samples on average). In cases in which data were acquired at higher sampling rates (for example because a participant’s computer monitor refresh rate exceeded 60 Hz), the data were down sampled to 60 Hz. Data were digitally smoothed using a second-order low-pass Butterworth filter with a cut-off frequency of 15 Hz. All data were stored for offline analysis using custom MATLAB R2020a (The MathWorks) scripts.

Movement trajectories were selected using an algorithm in which movement initiation was defined as the time at which the tangential velocity of the mouse first exceeded 0.5 cm/s and movement end was defined as the first time after peak velocity that tangential velocity fell below 0.5 cm/s, where peak velocity was defined as the fastest participants ever moved during the reach movement. For each trial we computed the angle between the line connecting the start position and the cursor position at peak velocity, and the line connecting the start position to the target. We determined the average reach angle, per subject during the last 50 trials of the baseline epoch and we subtracted this quantity from the reach angle measured on each trial.

### Model fitting

Smith et al. (3) outlined a method for mathematically modelling an iterative update of the states of the two proposed processes of short-term sensorimotor adaptation. Essentially, the model involves fitting four parameters: an error sensitivity and a retention parameter for both a fast and a slow process. The first parameter weighs the relative importance of recalling previous motor commands, which is interpreted as the retention factor. The second parameter is the sensitivity to error, which relates to the proportion of error that is corrected for trial-to-trial (1, 3, 4, 30). The two important assumptions in this model are that the error sensitivity is higher for the fast process compared with the slow process and that retention is stronger for the slow process compared with the fast process (3). Adaptation can be decomposed into a fast (**Eq. 1**) and a slow (**Eq. 2**) process, knowing that each state follows different learning dynamics. The two processes are summed together to produce the overall output x (**Eq. 3**). Error, denoted by e(n), arises on each trial n as the difference between the overall output xnet and the task parameter r (i.e., the degree of the rotation; **Eq.4**).

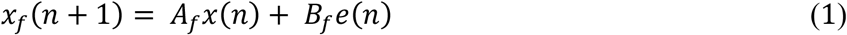

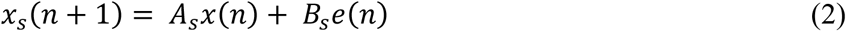

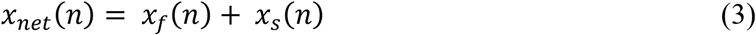

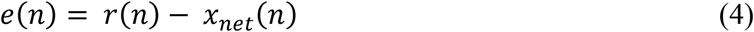

Linear inequality constraints were defined in order to apply to standard two-state model dynamics (31):

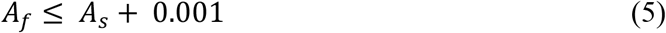

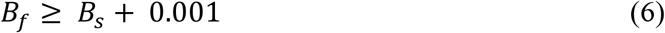

In order to approximate the four parameters (i.e., Af, As, Bf, and Bs), we fit the model to the behavioral data (using the function *fmincon* in MATLAB r2020a) by minimizing the squared difference between the estimated net output (xnet) of the model and the average participant reach angle, measured on each trial. According to the methods described in Albert and Shadmehr (31), we also included a mathematical formalization of visual error clamp trials and set breaks.

### Statistical Design

Pairwise comparisons were performed with nonparametric bootstrap hypothesis tests, as well as paired and unpaired t-tests. For statistical analyses that require multiple comparisons, we used the Holm-Bonferroni correction (32). Statistical tests were considered significant at p < 0.05. For all reported and depicted values, we report the mean and SEM.

## Results

Figure 2. shows the hand paths from one representative participant in the control group during both sessions one and two, as well as one representative participant per group in session two of the structural and gradual learning groups. During the baseline epoch (left column), these paths are relatively straight to the target. The representative participants were all adapting to an abrupt 30° CW rotation. During the early adaptation epoch (middle column) these movements were initially deviated in the CW direction, with a corrective movement at the end of the trajectory to bring the cursor to the target. In all three groups, participants adapted to the 30° CW rotation by the late adaptation epoch (right column), reducing their movement errors and resuming relatively straight hand paths to the target.

**Figure 2.**
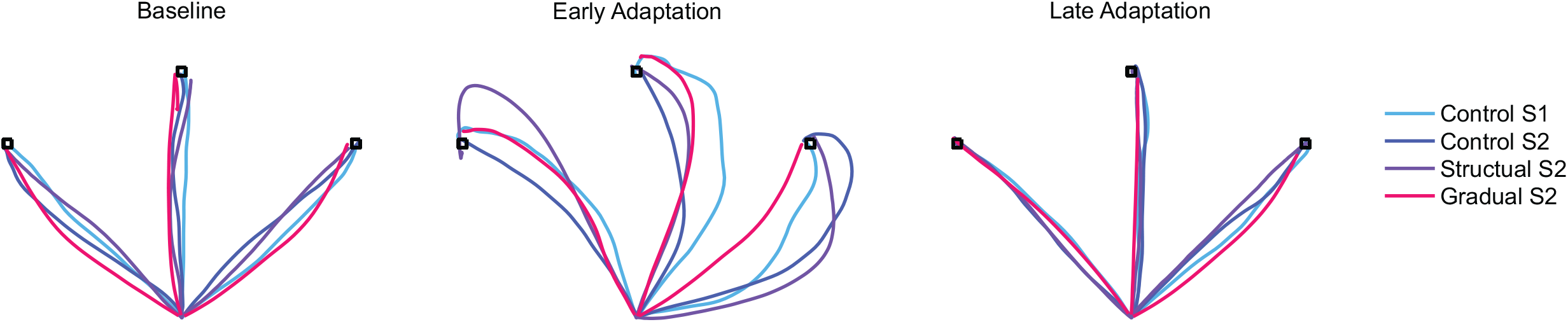
Hand trajectories from a representative participant in the control group during both session one (light blue) and two (dark blue), and one representative participant per group in session two of the structural (purple) and gradual (pink) learning groups. Baseline reaches were from the last three trials (from trial 68 to trial 70) during the baseline epoch. Early and late adaptation reaches were from the first (from trial 71 to 73) and last (from trial 368 to trial 370) three trials of the adaptation epoch, respectively. Participants saw a random ordering of the three possible targets (represented by the squares).

We used a kinematic behavioural measure to assess changes in performance. The primary outcome measure for the study was reach angle at peak velocity, which was measured as the angle between the straight line connecting the start position and the cursor position at peak velocity and the straight line connecting the start position to the target. The control group of participants adapted their movements to an abrupt 30° CW visuomotor rotation in both the first and second session. **Figure 3A** shows the angle at peak velocity for all trials in each session, averaged across participants in the control group. In both sessions, participants exhibited learning during the adaptation epoch, decay during the visual error clamp epoch, and a return towards baseline performance during the washout epoch. During the adaptation epoch we examined the learning at two different time points: early (first fifty trials during adaptation) and late (last fifty trials during adaptation; **Fig. 3B**). The mean angle in the early learning phase of the second session (*M* = 23.7, *SD* = 3.15) was reliably greater than in the first session [*M* = 19.9, *SD* = 4.6; paired *t*-test, *t*(19) = -6.2, *P* = 3.0e-06], indicating savings. We did not detect a reliable difference (*P* = 0.08) between sessions during late learning.

**Figure 3.**
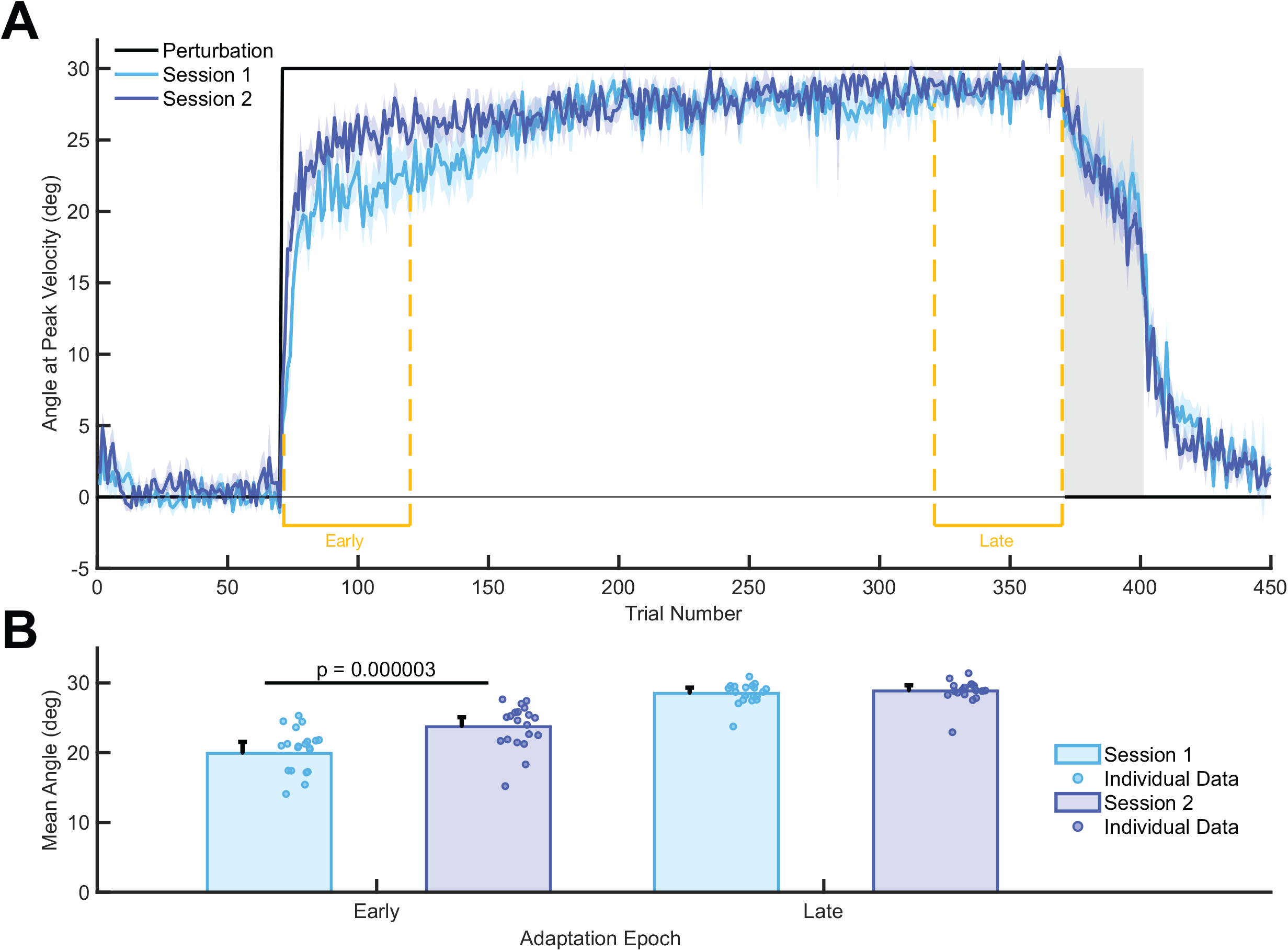
Control group. *A*: the average angle at peak velocity for all trials in *session 1* (light blue) and *session 2* (dark blue). The shaded region denotes ± SE. *B*: comparisons between *session 1* and *session 2* (dark blue) for the mean angle for the first 50 (early) and last 50 (late) trials of the adaptation epoch. Circles represent individual data.

A second group of participants was exposed to a gradual perturbation schedule during initial training. **Figure 4A** shows the angle at peak velocity for all trials in each session, averaged across participants in the gradual learning group. Participants exhibited learning during the adaptation epoch, decay during the visual error clamp epoch, and a return towards baseline performance during the washout epoch. A final group of participants was exposed to a series of brief exposures to large, random perturbations. Each participant in this group experienced a different set of randomly varying rotations. **Figure 4B** illustrates the angle at peak velocity for all trials in session one for four representative individual participants from the structural learning group. We observed two participants who demonstrated learning within each block of six trials, but who also appeared to have maintained a fraction of error throughout the adaptation epoch (**Fig. 4B, S2** and **S8**). In addition to a participant who adapted quickly to the randomly changing perturbation (**Fig. 4B, S18**), we observed a participant who qualitatively showed greater reduction of error in the latter half of the adaptation epoch, compared to the early half (**Fig. 4B, S20**).

**Figure 4.**
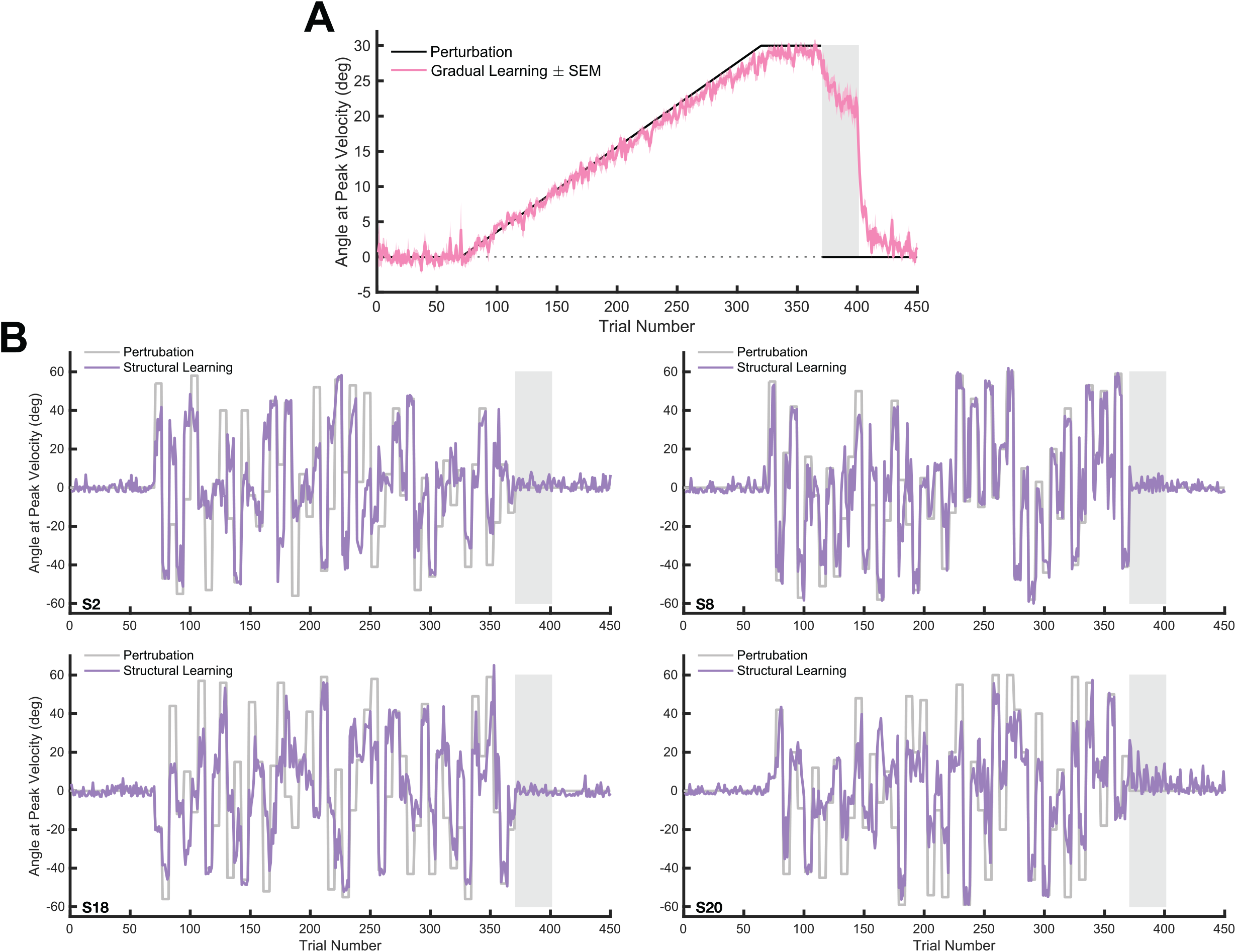
Gradual and structural learning groups. *A*: the average angle at peak velocity for all trials in *session 1* of the gradual learning group. The shaded region denotes ± SE. *B*: the data from four representative individual participants (S2, *top left*, S8 *top right*, S18 *bottom left*, S20 *bottom right*).

**Figure 5A** shows the angle at peak velocity averaged across participants for all trials in session one of the control group and session two of the structural and gradual learning groups. When comparing the model estimates of participants in the gradual and structural learning groups during the second session to the first session of the control group, we expected to see changes in error sensitivity that depended on the type of prior training participants experienced. To compare the changes in angle between the control, structural and gradual learning groups, we examined learning during the adaptation epoch at two different time points: early (first fifty trials during adaptation) and late (last fifty trials during adaptation; **Fig. 5B**). A one-way ANOVA revealed a significant effect of mean angle between the control, structural and gradual learning groups during early learning [*F*(2,57) = 14.4, *P* = 8.8 e-06].

**Figure 5.**
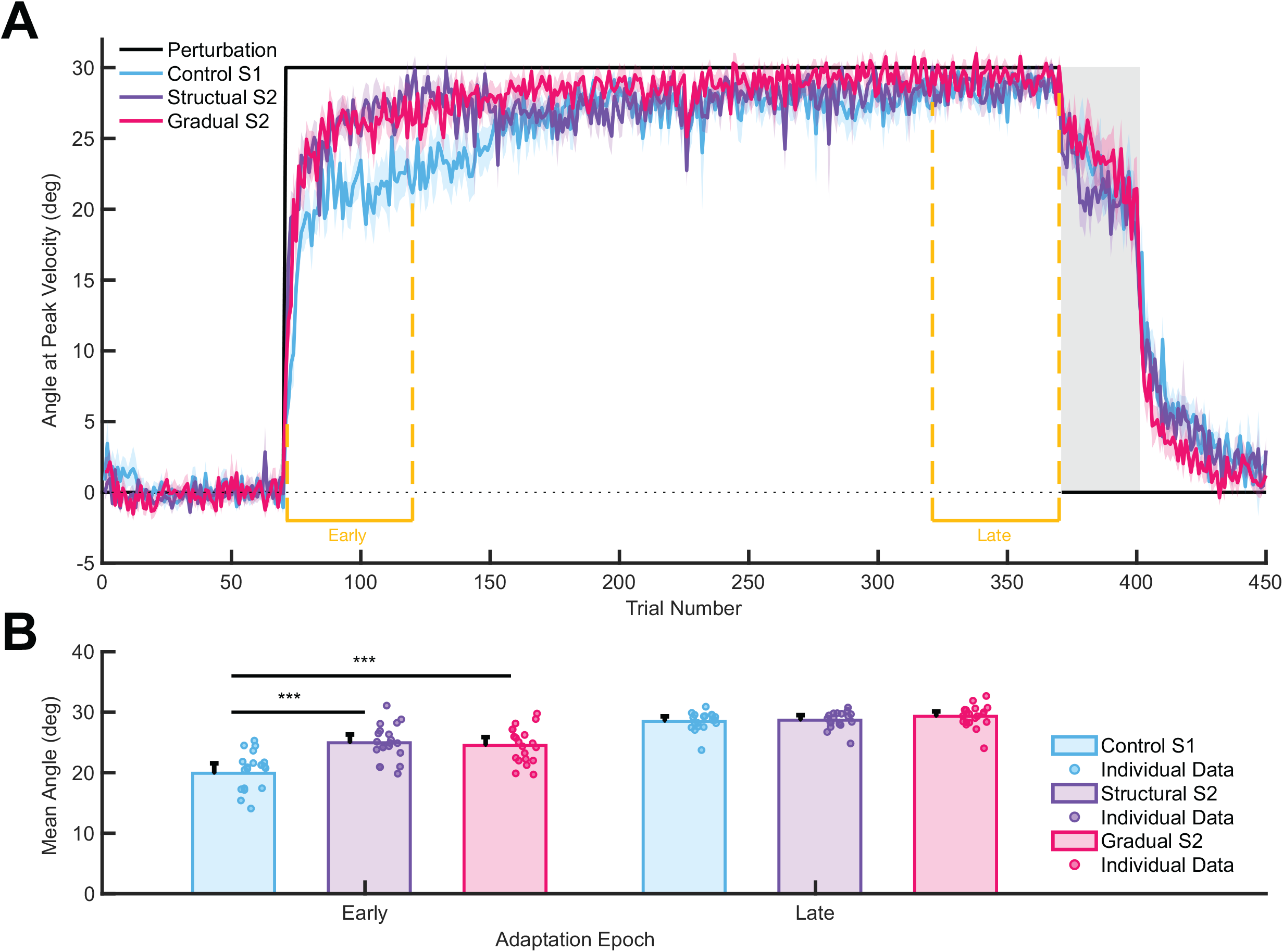
*A*: the average angle at peak velocity for all trials in *session 1* for the control group (light blue) and *session 2* for the structural (purple) and gradual (pink) learning groups. The shaded region denotes ± SE. *B*: comparisons between groups for the mean angle for the first fifty (early) and last 50 (late) trials of the adaptation epoch. Circles represent individual data.

*Post hoc* comparisons using Tukey HSD tests indicated that the mean angle for the structural learning group (*M* = 24.9, *SD* = 2.9, *P* = 2.9 e-05) and the gradual learning group (*M* = 24.5, *SD* = 2.9, *P* = 1.4 e-04) were reliably higher than the mean angle for the control group (*M* = 19.9, *SD* = 4.6). However, there was no reliable difference detected between the structural and gradual learning groups (*P* = 0.9). During late learning, we did not detect a reliable difference in mean angle among the groups (*P* = 0.2). Therefore, the structural and gradual learning groups demonstrated fast learning when countering an abrupt 30° CW rotation, as compared to session one of the control group. While the control group represented naive learners, the prior experience from session one for the structural and gradual learning groups is suggested to have facilitated the improved learning. Likewise, this was observed in the control group, in which participants experienced a repetition of an abrupt rotation and demonstrated savings during the second session.

Recent work suggests that error sensitivity in sensorimotor adaptation is likely not constant, but rather can vary depending on prior experience (13, 14, 16, 33). We modelled movement angle across each session with the state-space equations proposed by Smith et al. (3), and focused on changes in the retention and error sensitivity parameters. The main objective of this study was to compare the model parameters across groups learning to counter the abrupt 30° CW rotation. To do this, we used the bootstrap procedure previously reported by Coltman et al. (15). In this manner, we always fit the model to averaged group data for each resampled population (15, 31). The estimated posterior distributions of each of the four two-state model parameter values are depicted in **Fig. 6** for sessions one and two of the control group and session two of the gradual and structural learning groups. To determine whether the difference between the mean of each distribution was statistically reliable, we calculated the distribution of the differences in individual samples. The *insets* in **Fig. 6** show the distribution of differences found. **Table 1** shows the mean and standard deviation for each of the two-state parameters for each group.

**Table 1.**
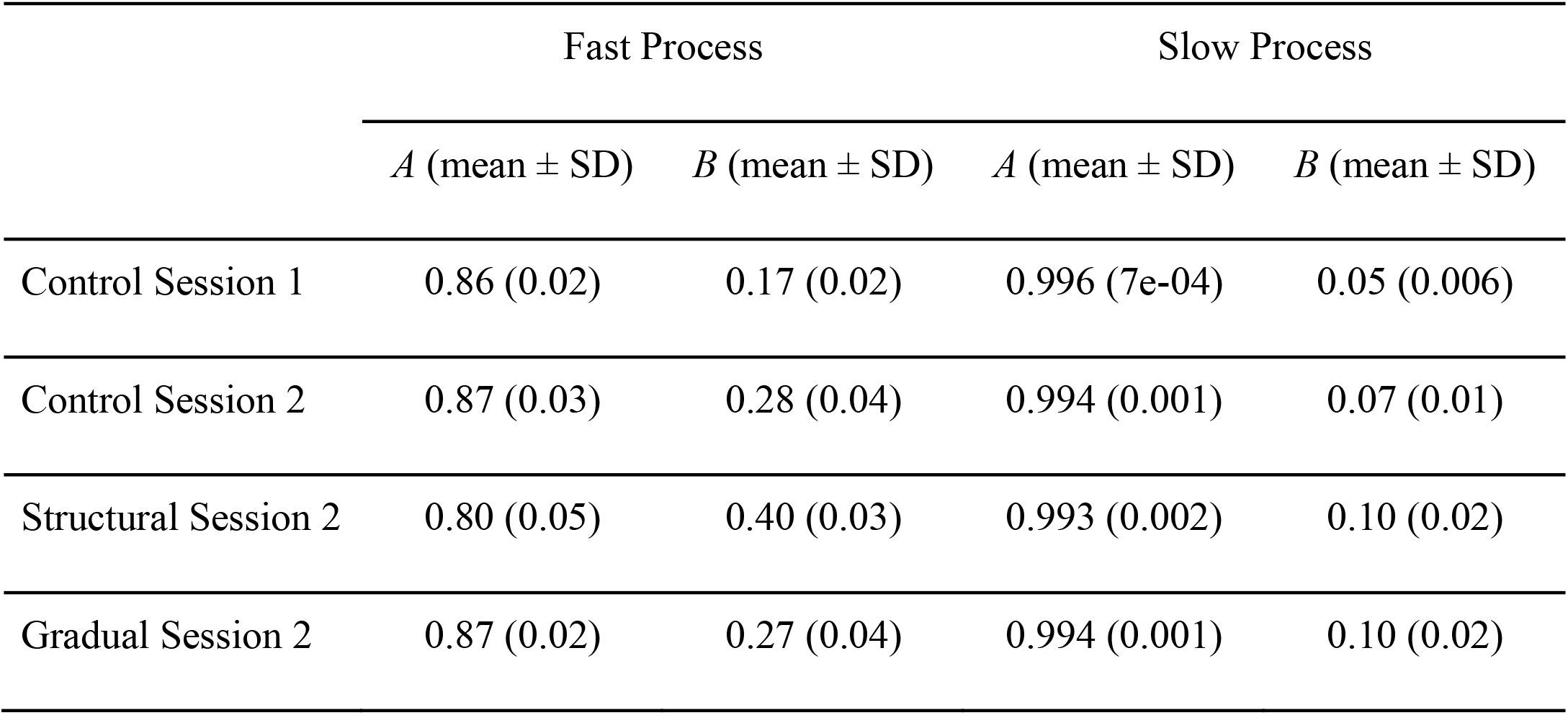
Two-state model parameters calculated from probability distribution

**Figure 6.**
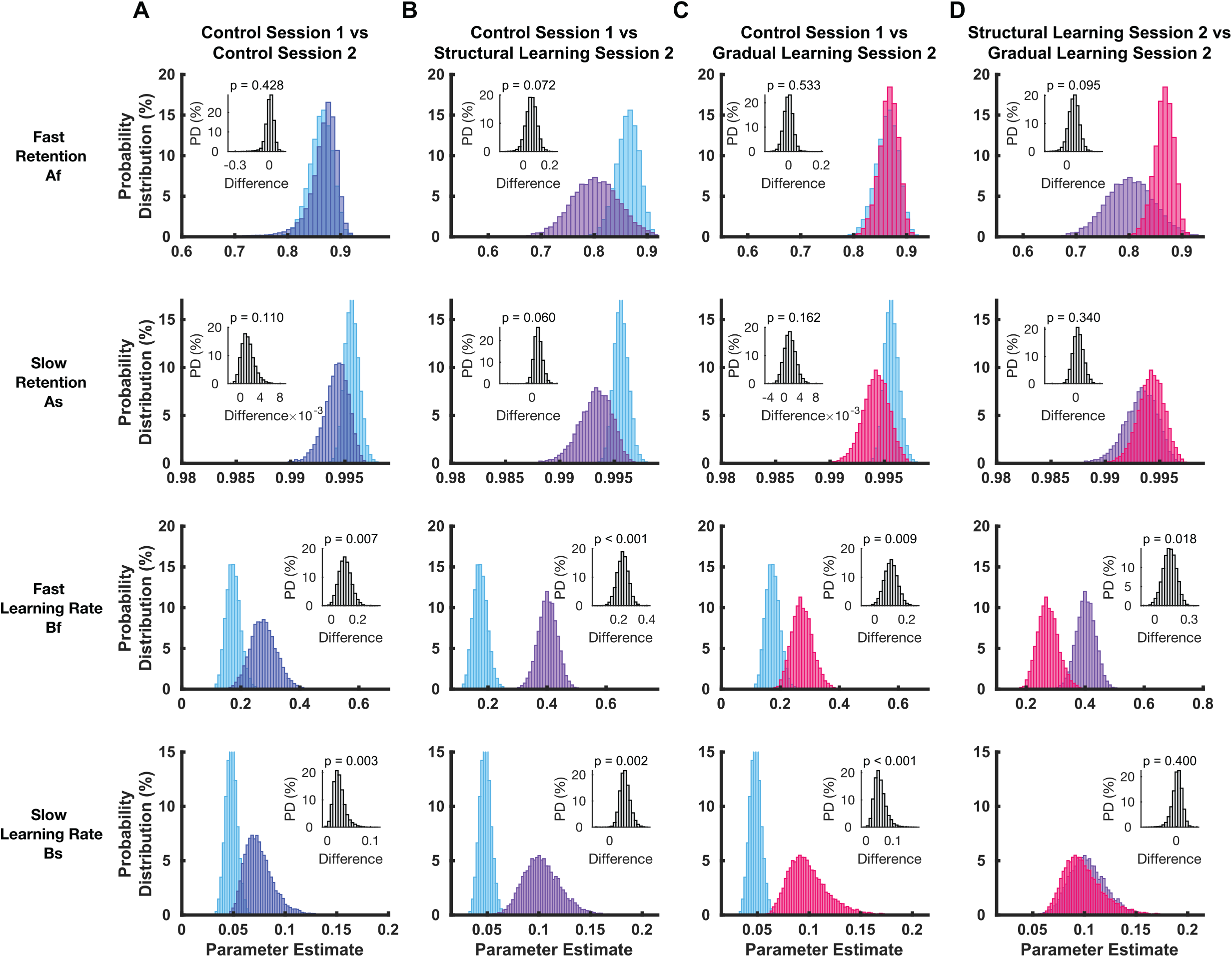
Probability distribution of the model parameters given the data. Light blue and dark blue represent *session 1* and *session 2* of the control group, respectively. Purple represents *session 2* of the structural learning group and Pink represents *session 2* of the gradual learning group. *Inset* represents the distribution of pairwise differences. The four model parameters of the two-state model are fast retention (A_f_), slow retention (A_s_), fast learning rate (B_f_), and slow learning rate (B_s_).

We first compared parameter estimates from session one and session two for the control group (**Fig. 6A**). Across all comparisons made between groups, we did not observe a reliable difference in the retention parameters for either the fast or slow process. When participants experienced repetition of the same abrupt rotation, we found a statistically reliable increase in the error sensitivity parameter for both the fast (B_*f*_, *P* = 0.007) and the slow (B_*s*_, *P* = 0.003) processes. Importantly, this comparison allowed us to demonstrate that our previous finding from a force field adaptation task (15) was replicated in a visuomotor rotation task. Therefore, this result suggests that both the fast and slow processes are responsive to a history of error and both contribute to savings.

Next, we compared parameter estimates from session one of the control group with session two of the structural learning group (**Fig. 6B**). Based on the theory of structural learning, thought to be essential to capturing the initial rapid phase of learning, Braun et al. (27) demonstrated that the benefit of knowing the underlying structure of a task is that it leads to facilitated adaptation. For this group we predicted that when later tested on an abrupt perturbation, only the fast process would be affected by the initial training, as compared to the control group. When overall learning is decomposed into a fast and slow state, the initial rapid phase of learning is dominated by the output of the fast process. Therefore, we assumed that such practice would influence the fast process. In addition to a statistically reliable increase in the error sensitivity parameter for the fast process (B_*f*_, *P* < 0.001), we also found a statistically reliable increase in the slow process error sensitivity (B_*s*_, *P* = 0.002).

Learning is believed to be more implicit in nature when a perturbation is gradually applied using small undetectable increases, so that participants never encounter large sensory prediction errors(22, 26, 34). By exposing a group of participants to a gradual perturbation schedule during initial training, we predicted that only the slow process would be influenced. When we compared the parameter estimates from session one of the control group with session two of the gradual learning group (**Fig. 6C**) we found the gradual learning group showed a statistically reliable increase in the error sensitivity parameter for both the fast (B_*f*_, *P* = 0.009) and the slow (B_*s*_, *P* = 2 e-04) processes.

Lastly, we compared parameter estimates between the structural and gradual learning groups during session two (**Fig. 6D**). Our goal was to use two different adaptation schedules thought to differentially affect fast and slow learning processes and test the idea that error sensitivity for each process would be independently modulated. We expected that the error sensitivity parameter for the fast process would be greater in the structural learning group compared to the gradual learning group, while the error sensitivity parameter for the slow process would be greater in the gradual learning group compared to the structural learning group. The only statistically reliable difference was in the error sensitivity parameter for the fast process that was larger for the structural learning group (B_*f*_, *P* = 0.02).

From the bootstrap distributions we calculated the mean value for each parameter for session one and session two of the control group, and session two of the structural and gradual learning groups separately. Using these mean estimated parameter values, we used the two-state model to simulate our experimental paradigm and generate simulated learning curves to visualize the time course of the estimated fast and slow processes, as well as the simulated overall output. **Figure 7** demonstrates that the simulated learning curves are qualitatively in good agreement with the measured behavioural data. The models explains 98 -99 % of the variance in angle over the course of learning (*control session 1*: R^2^ = 0.98, *P* = 2.2 e-04; *control session 2*: R^2^ = 0.99, *P* = 1.2 e-04; *structural session 2*: R^2^ = 0.98, *P* = 2.3 e-04; *gradual session 2*: R^2^ = 0.99, *P* = 1.6 e-04). The model effectively captures the initial improvement in learning during the adaptation epoch, the decay during the visual error clamp epoch, as well as the subsequent return towards baseline performance during the washout epoch.

**Figure 7.**
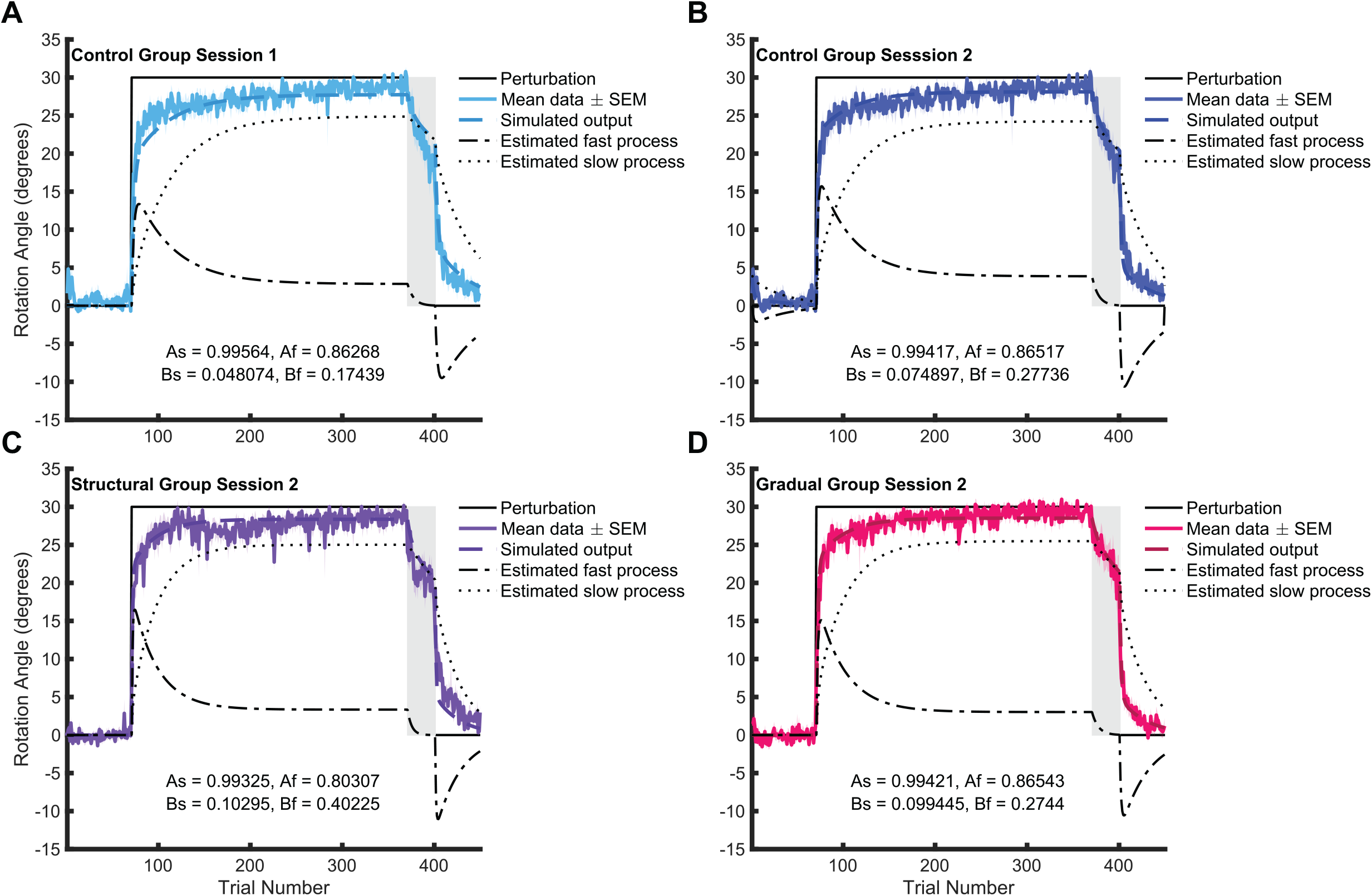
Model simulations. Parameter estimates for each session were based on the mean values from the bootstrap distributions shown in Fig. 6. The four parameters of the model are fast retention (Af), fast learning rate (Bf), slow retention (As), and slow learning rate (Bs).

## Discussion

The integration of different perturbation schedules and two-state modelling of measured behavioral data allowed us to test the role of prior experience on error sensitivity modulation during subsequent adaptation. The modelling of the data in turn describes adaptation as an interaction between error-sensitivity and retention. It has previously been shown in the context of force field learning that repetition of the same perturbation results in increased error sensitivity for both the fast and slow processes of adaptation (15). We substantiated this here by demonstrating that sensitivity to errors is similarly increased for both the fast and slow processes during the second session of a visuomotor rotation task. We found no reliable differences in the retention parameter across conditions and sessions.

The behavioural changes associated with savings suggest that some component of memory from the initial training must lead to the faster relearning, but what is remembered and recalled remains unclear. In the context of the present study, how the fast and slow processes individually contribute to savings, is not well known. To address this point, we used different perturbation schedules that relied on errors of different magnitudes to determine whether the underlying processes of adaptation could be independently manipulated, and whether an independent memory would subsequently be formed. We expected to see differences in error sensitivity depending on the type of prior training participants had received and therefore compared the model parameter estimates of participants in the gradual and structural learning groups to the first session of the control group, but we found that error sensitivity of both the fast and slow processes was increased for both groups. Such a result might suggest that sensitivity to error during visuomotor adaptation is modulated by abrupt, gradual and random perturbation schedules.

As an alternative account, savings has previously been explained by the retrieval of previous successful actions, reflecting the use of an explicit strategy (18-20, 23). Within the framework of a two-state model, this theory suggests that savings is driven purely by the fast process, without consideration of the contributions from the slow process (35). Several researchers have argued that explicit cognitive strategies can account for a significant amount of learning, particularly during the early phase of learning and relearning (36-38). The dissociation of learning into implicit and explicit learning processes often relies on the use of verbal aiming reports prior to reaching (18, 20, 23, 38). Recent findings, however, indicate that verbal aiming reports could lead to an overestimated explicit contribution to adaptation (21, 39). In fact, Leow et al. (21) demonstrated that the use of shortened preparation time, designed to prevent strategic re-aiming, resulted in the estimated implicit learning being larger than that which was obtained from verbal reports. Furthermore, Yin and Wei (22) provide supporting evidence that savings of motor adaptation is possible without forming or recalling a cognitive strategy with the use of a gradually introduced visuomotor rotation during initial learning. If savings is possible, with and without an explicit strategy being formed during initial learning and predominant measures of implicit and explicit processes may be confounding their mode of measurement, how reliable are the findings suggesting savings is driven exclusively by an explicit process?

The debate about the contributions of explicit/implicit and fast/slow processes to savings stems from a recent proposal that fast and slow processes reflect explicit and implicit learning mechanisms, respectively (35). Motor memory development is thought to be based on two components: recall, which involves retrieving past motor movements, and faster relearning, which involves increased sensitivity to errors (3, 40). By using the framework of a two-state model we focused on the dominant component of adaptation, which is sensory prediction error, and found that motor memory was associated with an increase in error sensitivity. In addition to sensory prediction error, there are other possible teaching cues that might drive adaptation. When researchers are focused on the more cognitive aspects of learning, exploring the use of an explicit strategy, the dominant component of adaptation may likely be the reinforcement of successful actions. Future research may shed light on this debate by probing both proposed methods of dual processing simultaneously during learning, and by assessing more directly their shared features.

Another long-standing question is how quickly implicit changes in learning emerge. Huberdeau and colleagues (20) demonstrated that learning of an abrupt perturbation with only a few trials is sufficient to cause savings via the explicit process, based on the belief that the fast learning is too short for implicit learning to take its full effect. Ruttle et al. (41) however recently confronted the long standing notion that implicit learning is slowly developing, typically unfolding over tens of trials. By observing changes in both internal models and state estimates of limb position as a characterization of implicit learning, they found that after only one to three perturbed training trials participants had changes in both reach aftereffects and a shift in hand localization. Taking this into account, it seems possible that the 6-trial repetition used in the structural learning task, aiming to influence the fast process, may have simultaneously influenced the slow process. For that reason, it is possible that a common component of all three perturbation schedules used during initial training was that the slow process accounted for a significant portion of the error reduction.

Albert et al. (33) recently investigated the persistence of residual errors during motor adaptation in the context of implicit and explicit learning systems. Of importance to the present study, they propose that it is the implicit learning system which maintains a history for prior errors. Our results are consistent with this hypothesis that it is the implicit process that stored some component of prior training. Given the suggestion that the history of errors is stored by only one of the two proposed underlying processes, this finding would be lost if learning behavior was represented using a single-state model. Nevertheless, one may ask whether a two-state model was necessary to represent learning in the behavioral tasks tested in the present study. To address this, we calculated AIC values for both single-and two-state models fits to the behavioral data. We used the data associated with the four sessions of abrupt rotations (i.e., the first and second sessions of the control group, and the second session of the gradual and structural learning groups) and for each we estimated the overall output based on a single-state model and separately using a two-state model. Based on the model with the lowest AIC value, in all four cases the best-fit model describing the measured behavioural data was the two-state model. As a follow-up to our initial question, we would further suggest that the stored memory is accessible to both processes during subsequent learning. As it pertains to our findings, we would argue that while the fast process may not maintain a history or errors, it does have access to this information in subsequent learning as evident by the increased error-sensitivity for the fast process during testing in all groups.

Alternatively, while the experimental design and two-state model, used in the present study, account well for the results of savings, recent work looking at evoked recovery (42) posits that memory formation is related to the storing of information about the dynamical and sensory features of the environment is related to the context with which it is associated. Understanding how contextual inference can be related to and accessed by each process of the two-state model can shed light on future discussions about multiple processes underlying motor learning.

## Acknowledgements

Canadian Institutes of Health Research (CIHR) and the Natural Sciences and Engineering Council of Canada (NSERC).

## References

1. Donchin O, Francis JT, and Shadmehr R. Quantifying generalization from trial-by-trial behavior of adaptive systems that learn with basis functions: theory and experiments in human motor control. Journal of Neuroscience 23: 9032–9045, 2003.

2. Miall RC, and Wolpert DM. Forward models for physiological motor control. Neural networks 9: 1265–1279, 1996.

3. Smith MA, Ghazizadeh A, and Shadmehr R. Interacting adaptive processes with different timescales underlie short-term motor learning. PLoS Biol 4: e179, 2006.

4. Thoroughman KA, and Shadmehr R. Learning of action through adaptive combination of motor primitives. Nature 407: 742–747, 2000.

5. Wolpert DM, and Kawato M. Multiple paired forward and inverse models for motor control. Neural networks 11: 1317–1329, 1998.

6. Kording KP, Tenenbaum JB, and Shadmehr R. The dynamics of memory as a consequence of optimal adaptation to a changing body. Nature neuroscience 10: 779–786, 2007.

7. Lee J-Y, and Schweighofer N. Dual adaptation supports a parallel architecture of motor memory. Journal of Neuroscience 29: 10396–10404, 2009.

8. Coltman SK, and Gribble PL. Time course of changes in the long-latency feedback response parallels the fast process of short-term motor adaptation. J Neurophysiol 124: 388–399, 2020.

9. Kim S, Ogawa K, Lv J, Schweighofer N, and Imamizu H. Neural substrates related to motor memory with multiple timescales in sensorimotor adaptation. PLoS biology 13: e1002312, 2015.

10. Sarwary AM, Wischnewski M, Schutter DJ, Selen LP, and Medendorp WP. Corticospinal correlates of fast and slow adaptive processes in motor learning. Journal of neurophysiology 120: 2011–2019, 2018.

11. Ebbinghaus H. Memory: A Contribution to Experimental Psychology, eds. HA Ruger and CE Bussenius (Trans)(Dover Publications, New York, 1964)(Original Work Published 1885) 1913.

12. van Beers RJ. Motor learning is optimally tuned to the properties of motor noise. Neuron 63: 406–417, 2009.

13. Wei K, and Kording K. Relevance of error: what drives motor adaptation? Journal of neurophysiology 101: 655–664, 2009.

14. Marko MK, Haith AM, Harran MD, and Shadmehr R. Sensitivity to prediction error in reach adaptation. Journal of neurophysiology 108: 1752–1763, 2012.

15. Coltman SK, Cashaback JG, and Gribble PL. Both fast and slow learning processes contribute to savings following sensorimotor adaptation. Journal of neurophysiology 121: 1575–1583, 2019.

16. Herzfeld DJ, Vaswani PA, Marko MK, and Shadmehr R. A memory of errors in sensorimotor learning. Science 345: 1349–1353, 2014.

17. Leow L-A, De Rugy A, Marinovic W, Riek S, and Carroll TJ. Savings for visuomotor adaptation require prior history of error, not prior repetition of successful actions. Journal of neurophysiology 116: 1603–1614, 2016.

18. Avraham G, Morehead JR, Kim HE, and Ivry RB. Reexposure to a sensorimotor perturbation produces opposite effects on explicit and implicit learning processes. PLoS biology 19: e3001147, 2021.

19. Huang VS, Haith A, Mazzoni P, and Krakauer JW. Rethinking motor learning and savings in adaptation paradigms: model-free memory for successful actions combines with internal models. Neuron 70: 787–801, 2011.

20. Huberdeau DM, Haith AM, and Krakauer JW. Formation of a long-term memory for visuomotor adaptation following only a few trials of practice. Journal of neurophysiology 114: 969–977, 2015.

21. Leow L-A, Gunn R, Marinovic W, and Carroll TJ. Estimating the implicit component of visuomotor rotation learning by constraining movement preparation time. Journal of neurophysiology 118: 666–676, 2017.

22. Yin C, and Wei K. Savings in sensorimotor adaptation without an explicit strategy. Journal of neurophysiology 123: 1180–1192, 2020.

23. Morehead JR, Qasim SE, Crossley MJ, and Ivry R. Savings upon re-aiming in visuomotor adaptation. Journal of neuroscience 35: 14386–14396, 2015.

24. Hanajima R, Shadmehr R, Ohminami S, Tsutsumi R, Shirota Y, Shimizu T, Tanaka N, Terao Y, Tsuji S, and Ugawa Y. Modulation of error-sensitivity during a prism adaptation task in people with cerebellar degeneration. Journal of neurophysiology 114: 2460–2471, 2015.

25. Kim HE, Morehead JR, Parvin DE, Moazzezi R, and Ivry RB. Invariant errors reveal limitations in motor correction rather than constraints on error sensitivity. Communications Biology 1: 1–7, 2018.

26. Orban de Xivry J-J, and Lef. vre P. Formation of model-free motor memories during motor adaptation depends on perturbation schedule. Journal of neurophysiology 113: 2733–2741, 2015.

27. Braun DA, Aertsen A, Wolpert DM, and Mehring C. Motor task variation induces structural learning. Current Biology 19: 352–357, 2009.

28. Braun DA, Mehring C, and Wolpert DM. Structure learning in action. Behavioural brain research 206: 157–165, 2010.

29. Bond KM, and Taylor JA. Structural learning in a visuomotor adaptation task is explicitly accessible. Eneuro 4: 2017.

30. Scheidt RA, Dingwell JB, and Mussa-Ivaldi FA. Learning to move amid uncertainty. Journal of neurophysiology 86: 971–985, 2001.

31. Albert ST, and Shadmehr R. Estimating properties of the fast and slow adaptive processes during sensorimotor adaptation. Journal of Neurophysiology 119: 1367–1393, 2018.

32. Holm S. A simple sequentially rejective multiple test procedure. Scandinavian journal of statistics 65–70, 1979.

33. Albert ST, Jang J, Sheahan HR, Teunissen L, Vandevoorde K, Herzfeld DJ, and Shadmehr R. An implicit memory of errors limits human sensorimotor adaptation. Nature Human Behaviour 2021.

34. Criscimagna-Hemminger SE, Bastian AJ, and Shadmehr R. Size of error affects cerebellar contributions to motor learning. Journal of neurophysiology 103: 2275–2284, 2010.

35. McDougle SD, Bond KM, and Taylor JA. Explicit and implicit processes constitute the fast and slow processes of sensorimotor learning. Journal of Neuroscience 35: 9568–9579, 2015.

36. Mazzoni P, and Krakauer JW. An implicit plan overrides an explicit strategy during visuomotor adaptation. Journal of neuroscience 26: 3642–3645, 2006.

37. Taylor JA, and Ivry RB. Flexible cognitive strategies during motor learning. PLoS computational biology 7: e1001096, 2011.

38. Taylor JA, Krakauer JW, and Ivry RB. Explicit and implicit contributions to learning in a sensorimotor adaptation task. Journal of Neuroscience 34: 3023–3032, 2014.

39. de Brouwer AJ, Albaghdadi M, Flanagan JR, and Gallivan JP. Using gaze behavior to parcellate the explicit and implicit contributions to visuomotor learning. Journal of neurophysiology 120: 1602–1615, 2018.

40. Mawase F, Bar-Haim S, and Shmuelof L. Formation of Long-term Locomotor memories is associated with functional connectivity changes in the cerebellar–thalamic–cortical network. Journal of Neuroscience 37: 349–361, 2017.

41. Ruttle JE, Marius’t Hart B, and Henriques DY. Implicit motor learning within three trials. Scientific Reports 11: 1–11, 2021.

42. Heald J, Lengyel M, and Wolpert D. Contextual inference underlies the learning of sensorimotor repertoires. bioRxiv 2020.

